# Computational mechanisms of osmoregulation: a reinforcement learning model for sodium appetite

**DOI:** 10.1101/2021.04.20.440596

**Authors:** Yuuki Uchida, Takatoshi Hikida, Yuichi Yamashita

## Abstract

Homeostatic control with oral nutrient intake is a vital complex system involving the orderly interactions between the external and internal senses, behavioral control, and reward learning. Sodium appetite is a representative system and has been intensively investigated in animal models of homeostatic systems and oral nutrient intake. However, the system-level mechanisms for regulating sodium intake behavior and homeostatic control remain unclear.

In the current study, we attempted to provide a mechanistic understanding of sodium appetite behavior by using a computational model, the homeostatic reinforcement learning model, in which homeostatic behaviors are interpreted as reinforcement learning processes. Through simulation experiments, we confirmed that our homeostatic reinforcement learning model successfully reproduced homeostatic behaviors by regulating sodium appetite. These behaviors include the approach and avoidance behaviors to sodium according to the internal states of individuals. In addition, based on the assumption that the sense of taste is a predictor of changes in the internal state, the homeostatic reinforcement learning model successfully reproduced the previous paradoxical observations of the intragastric infusion test, which cannot be explained by the classical drive reduction theory. Moreover, we extended the homeostatic reinforcement learning model to multi-modal data, and successfully reproduced the behavioral tests in which water and sodium appetite were mediated by each other. Finally, through an experimental simulation of chemical manipulation in a specific neural population in the brain stem, we proposed a testable hypothesis for the function of neural circuits involving sodium appetite behavior.

The study results support the idea that osmoregulation via sodium appetitive behavior can be understood as a reinforcement learning process and provide a mechanistic explanation for the underlying neural mechanisms of sodium appetite and homeostatic behavior.

**Author Summary:** The taste of high-concentration saltwater is rewarding during sodium depletion, while it is aversive in a sodium sufficient state. This “sodium appetite” is a clear manifestation of homeostasis maintenance and proper action selection in animals, reflecting the internal environment. To reveal the computational mechanism of this property, we applied a machine learning model, in which homeostatic stability is a reward and the goal is to maximize the sum of the reward, and simulated animal behavioral experiments. The results suggest that the mechanism of sodium-appetite behavior is based on the machine learning model. Specifically, by replicating the results of neural circuit manipulation, which controls sodium appetite, an algorithm in which the function of a neural population affects sodium appetite behaviors was proposed. Our results provide a fundamental computational model for a mechanism by a function of a neural cell type to regulate animal behavior. More generally, this study can be fundamental to understanding the computational process of decision making reflecting the internal environment.

## Introduction

Homeostatic control systems with oral nutrient intake are vital to sustain life. The system is quite complex, in which orderly interactions between external and internal senses, behavioral control, and reward learning are essential [1, 2]. Their failure to properly develop or maintain precisely results in several disease symptoms. For example, taste disorders may cause collapse of homeostasis and nutrient disorders [3-5].

Sodium appetite is a representative system that has been intensively investigated in animal models of the homeostatic systems that coordinate oral nutrient intake [6-11]. For example, at the behavioral level, it is known that preference for salty taste changes depending on the internal sodium state. That is, when an animal is sodium-depleted after adrenalectomy [12], administration of furosemide, or low-sodium food [9, 10], it exhibits a positive sodium appetite, which means that sodium intake works as a reward. On the other hand, when an animal is not deficient in sodium, it will exhibit a negative sodium appetite, and sodium intake works as a punishment [13, 14]. This property is related to various factors, and not only to avoidance of harmful foods or the consumption of essential nutrients. Rather, it is a complex phenomenon in which the reward value of a taste fluctuates, reflecting the internal states of animals.

At the physiological level, multiple hormones are involved in controlling sodium appetite. For instance, adrenalectomized rats, which have difficulty secreting aldosterone, show increased sodium appetite [6]. In relation to this, sodium deficiency increases the level of angiotensin II and stimulates the secretion of aldosterone [15].

Furthermore, the pacemaker-like firing of aldosterone-sensing neurons in the nucleus of the solitary tract (NTS^HSD2^ neurons) has been observed in sodium-deficient animals [16].

At the neural substrate level, it is known that several neuronal nuclei (e.g., limbic system, pons, and basal ganglia) control sodium appetite. For example, activation of the ventral tegmental area (VTA) dopaminergic neurons, which show robust correlations with reward systems, decreases salt intake [17]. Dopaminergic neurons in the midbrain may encode appetitive properties of sodium [18]. Some excitatory neurons in the pre-locus coeruleus decrease sodium appetite [10]. Conversely, the activation of neurons in the subfornical organ increases sodium appetite [9].

Despite these findings, the system-level mechanisms for regulating sodium intake behavior and homeostatic controls remain unclear. The drive reduction theory [19] is a classical depiction that has provided a basis for the nature of sodium appetite behavior; that is, the discrepancy from the optimal state drives behavior to reduce the discrepancy. However, some important biological observations cannot be explained by the drive reduction theory. For example, direct infusion of sodium into the stomach, not through the mouth, does not reduce sodium appetite for several minutes [2], even though the internal sodium state may approach the optimal level.

To address this issue, here we propose a computational model to provide a mechanistic understanding of sodium appetite, the homeostatic reinforcement learning (HRL) model, according to which homeostatic behaviors are interpreted as reinforcement learning processes [2, 20, 21]. In the HRL model, changes in the internal state approaching the ideal states were regarded as rewards, while changes in the internal state moving away from the ideal state were considered as punishment. Based on this idea, the values of the optimal behavior to maintain internal states were acquired through an incremental learning process. In addition, by treating the taste modality and the actual change of internal state separately, the HRL model provides explanations about the mechanisms that integrate taste, behavior, and maintenance of inner environments.

The HRL model was originally proposed to explain homeostatic controls for body temperature, internal water state [2], and pathological mechanisms of cocaine addiction [21]. This study is the first attempt to use the HRL model to explain sodium appetite behavior. In addition, although the HRL model can handle multi-dimensional internal states, previous studies have utilized it to treat one-dimensional changes in the internal state [2, 20, 21]. However, sodium appetite is a complex process in which the interactions and homeostatic balance between the preference for water and sodium are involved. Therefore, in the current study, we extended the HRL model to a multi-dimensional model by simulating two internal states of water and sodium. Based on this extension, we attempted to provide a mechanistic understanding of previous findings using experiments with dynamic interactions between the internal states of water and sodium. Finally, by simulating an experimental manipulation of the neural activity of a particular brain nucleus known to be involved in sodium appetitive behavior, we provide a hypothesis for the function of a specific neural population in the control of sodium appetite.

## Results

### Model overview

In the current study, sodium appetitive behavior was modeled using the HRL model [2, 20, 21]. This model is based on the assumption that homeostasis is an RL process, in which the minimization of deviations of internal states from an optimal level (i.e., homeostasis) is treated as a computation for maximizing the sum of rewards. In the HRL model, a multi-dimensional metric space in which each dimension represents an internal state (such as body temperature, blood glucose density, water state, and sodium level) was defined as a “homeostatic space”. In this homeostatic space, the drive function *D(H*_*t*_) was defined as the distance between the internal state of the *i*-th component (e.g., water or sodium) at time *t, H*_*i,t*_, and the ideal internal state *H\**_*i*_:

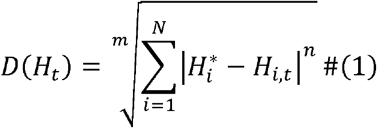

Where *m* and *n* are free parameters to define the distance, *and N* is the total number of dimensions for internal states (e.g., water, sodium, etc.). When the internal state approaches the ideal state, the value of the drive function should be reduced. Based on this drive function, the reward *r* is determined as a change in the values of the drive function from time *t* to time *t+1*. Specifically, to implement nutrient intake, the internal state at time *t+1* should contain the amount of nutrient intake at time *t*, defined as *K*_*t*_ :

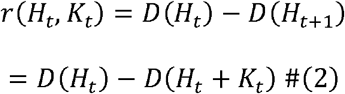

As described later, in the HRL model, the intake of taste stimuli (*K^*) can be modeled as a predictor of the actual nutrient intake (*K*). Under this assumption, the reward was calculated as follows:

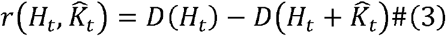

The RL process was modeled as a Rescorla-Wagner model. In this model, the values of action *a* (e.g., sodium intake, do nothing…), *Q(a)*, are renewed based on the reward prediction error:

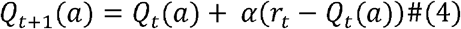

Where, α is the learning rate.

### Experiment 1: Sodium homeostasis according to the HRL

First, we confirmed that the homeostatic control of the internal states of sodium can be replicated using the framework of the HRL model. In this experiment, model mice were able to choose to either perform *saltwater intake* or *do nothing* (Fig 1B). The action values of intake and do nothing were both set to 0. Therefore, the initial action selection was random (P(intake) = 0.5) (Fig 1C). After several random choices of sodium intake, the action value of intake was reinforced, and the internal state approached the ideal state. In approximately 20 trials, P(intake) was nearly 1, and the internal state rapidly reached the ideal state. When the internal state exceeded the ideal state after approximately 80 trials, sodium intake became a punishment, and the action value of *do nothing* increased. After several trials of *do nothing*, at around trial number 140, due to the decay assumed to be a natural loss of internal sodium (see Method for more details), the internal state became lower than the ideal state. Therefore, the action value of sodium intake increased again. Through the repetitions of this cycle, the model successfully achieved homeostatic control of the internal states of sodium (Fig 1C).

**Fig 1.**
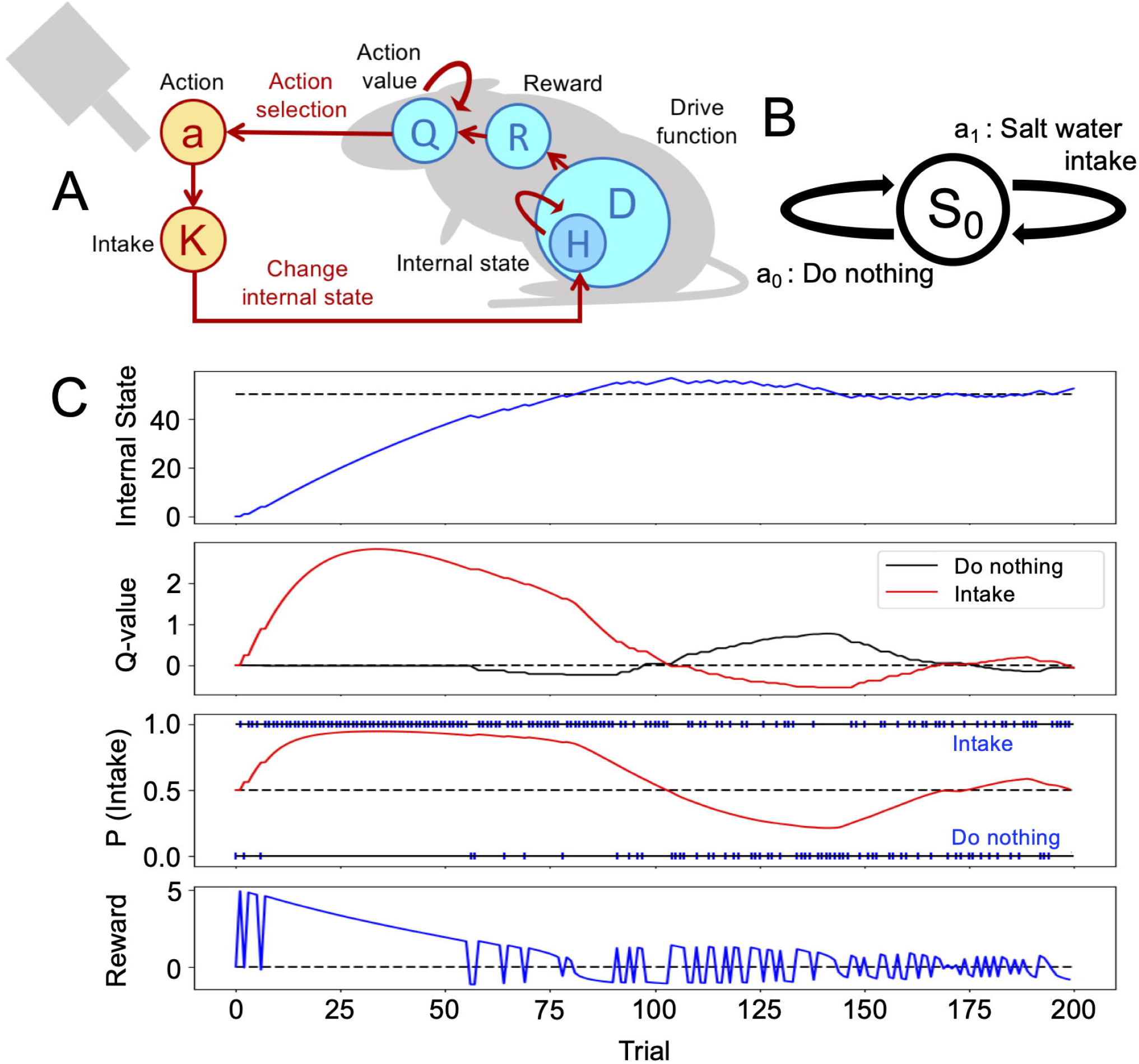
Homeostatic behavior according to the homeostatic reinforcement learning (HRL) model. (A) Schematic drawing of the computational process of the HRL model. (B) Definition of a state and two actions in experiments 1 and 2. (C) Example of homeostatic behavior. Changes in internal sodium state, the value of each action (Q-value), probability of sodium intake (P(Intake)), and magnitude of reward over time were plotted. The dotted line in the panel of the internal state indicates the ideal point (H = 50) of sodium taste. In the panel of the probability of sodium intake (P(Intake)), the blue marks on axis P = 1.0 represent ‘Intake behavior’, and those on axis P = 0.0 indicate ‘Do nothing’. At the beginning of the simulation, the internal sodium state and Q-values for each action were set to 0. After several random selections of action, the sodium Q-value of intake was increased, and the internal sodium state quickly reached the ideal point, maintaining homeostatically-regulated behaviors.

### Experiment-2: Sense of taste as a predictor of changes in internal states

In the homeostatic control of sodium states, as well as in the monitoring of the internal states, the sense of taste may play an important role. In the HRL model, this assumption can be tested by implementing this sensory modality as a predictor of changes in the internal states induced by nutrient intake [2]. In this study, we simulated an intragastric infusion test [10]. In the intragastric infusion test, there were three groups of animals: (1) a control group of sodium-depleted mice, (2) a NaCl-IG group of sodium-depleted animals treated with intragastric infusion of saltwater before the test, and (3) a NaCl-oral group with sodium-depleted mice stimulated with sodium (salty taste) during the test (Fig 2B). The definitions of the states and actions were the same as in the previous experiment (Fig 1B). At the beginning of the intragastric infusion test, the internal states for the control group and NaCl-oral group were set at *H*_*t*_ = 0, while for the NaCl-IG group, the internal state was set at *H*_*t*_ = *H\**/2, corresponding to sodium partially supplied through a gastric infusion. Each animal model was tested in 100 trials, with a total duration of 600 s in the actual experiments. During the 100 trials, the model animals of the NaCl-oral group were assumed to have constant salty taste stimulation (Fig 2A).

**Fig 2.**
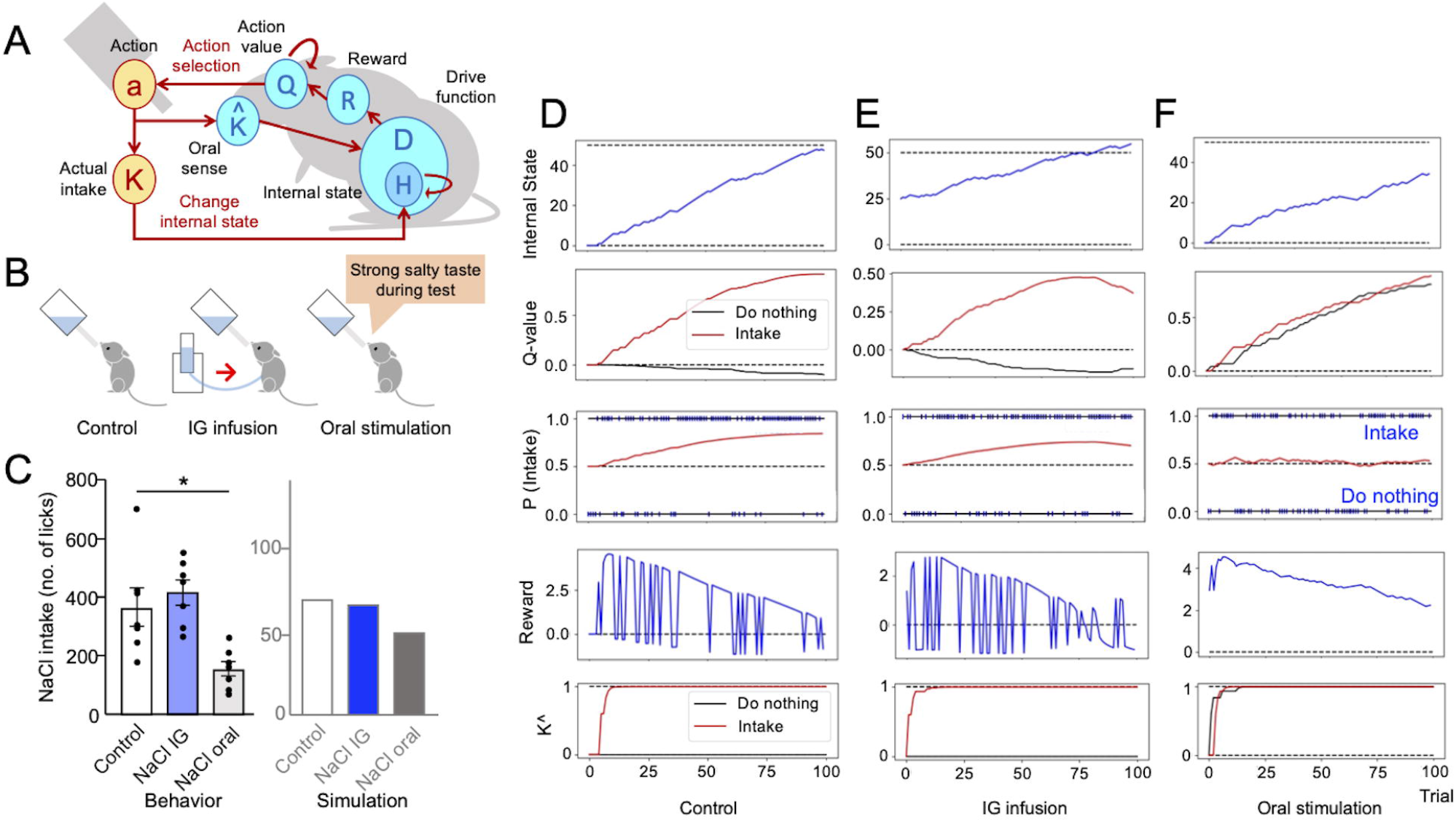
Intragastric and oral stimulation tests were explained using the sense of taste as a predictor of changes in the internal state. (A) Schematic illustration of experiments 2 and 3: Taste perception (K^) was a predictor of an increase in the internal state. (B) Three groups of behavioral tests. (C) Comparison of results of the behavioral experiments [10] and the computational model. The intragastric infusion did not change sodium intake, while oral stimulation decreased the intake. (D-F) Transitions of the simulation. (D) The control group shows an increased Q-value of intake. (E) The congenial increase of the Q-value is seen in the NaCl-IG group. (F) The values of ‘Do nothing’ and ‘Intake’ are reinforced in the NaCl-oral group.

As a result of the simulation, there was no significant difference in the total numbers of NaCl intakes between the control and NaCl-IG groups (Fig 2C). On the other hand, the total intake of the NaCl-oral group was clearly lower than that of the control and NaCl-IG groups. These trends were similar to those observed in animal experiments [10] (Fig 2C).

To provide a mechanistic overview of these results, the transitions of each variable during the experiment are plotted in Fig. 2D-2F. In the NaCl-IG group, even though sodium was partly supplied through gastric infusion, an increase in the action value of sodium intake led to an increase in sodium intake behavior (Fig 2E), resulting in the number of intakes not changing significantly compared with the control group (Fig 2C).

On the other hand, in the NaCl-oral group, the action value of do nothing was reinforced (Fig 2F) based on the expectation of an increase in the internal sodium state (K^) due to the continuous application of the salty stimulus (sodium) during the test. There were no clear differences between the action values of salt intake and do nothing (Fig 2F), and the number of intakes of the group that received NaCl as drinking water was lower than that of the control group (Fig 2C).

### Experiment 3: Multi-dimensional HRL

In this experiment, in order to describe the dynamic interaction between the internal states of water and sodium, we extended the HRL model to multiple dimensions [2]. The HRL model with multi-dimensional internal states (water state and sodium state) was assessed with a two-bottle preference test, which is a behavioral procedure used to compare the preference toward contents in two bottles (Fig 3B). In the two-bottle preference test, the model mice were able to either perform “saltwater intake,” “water intake”, or “do nothing” (Fig 3A). In this experiment, there were four groups of animals: a control group with fulfilled initial internal states, a sodium-depleted group, a water-depleted group, and a water/salt-depleted group.

**Fig 3.**
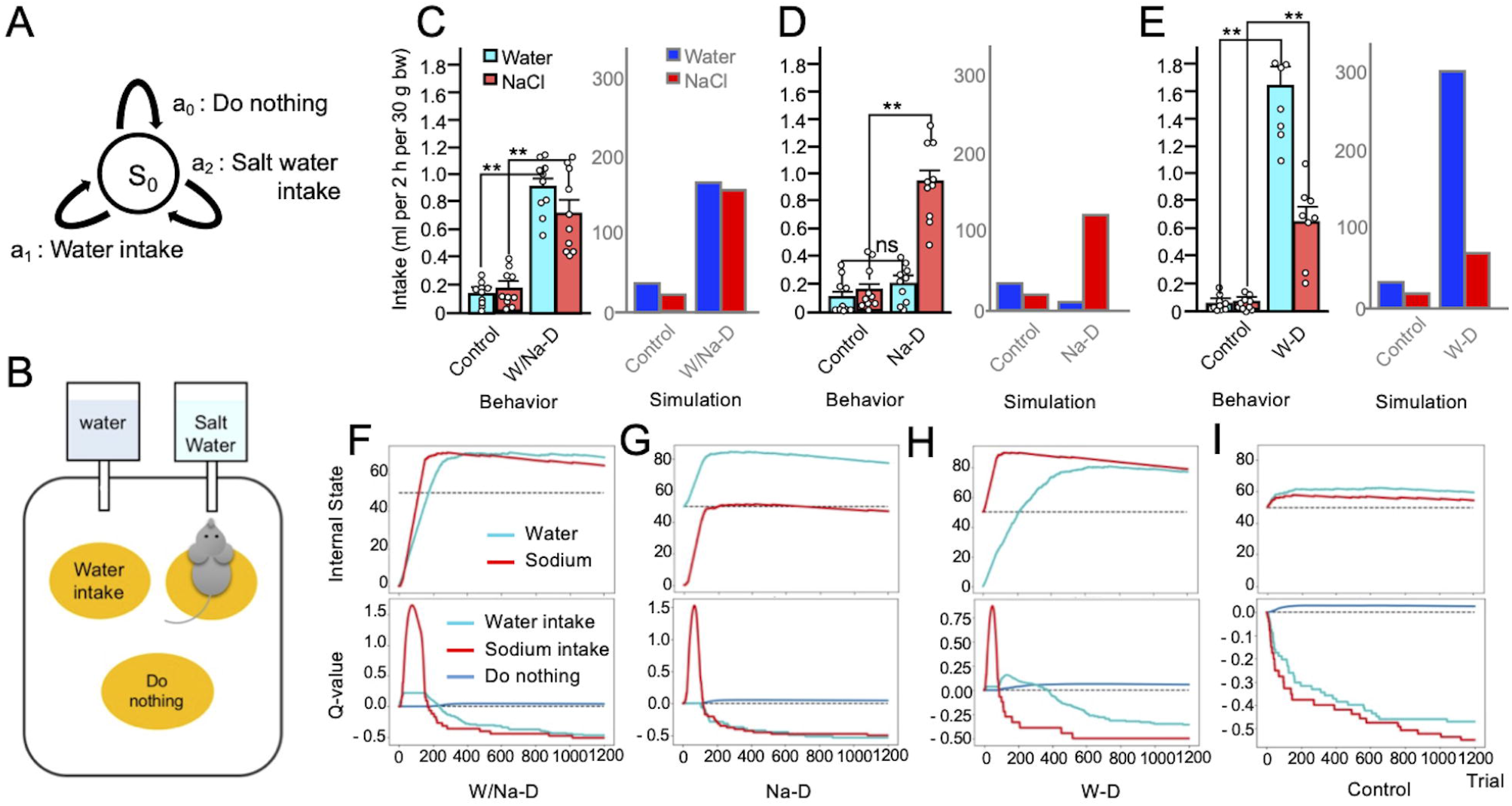
Multi-dimensional homeostatic reinforcement learning (HRL) model as a suitable explanation of the result of the two-bottle preference test. (A) Definition of a state and three actions in experiments 3 and 4. (B) Schematic drawing of experiment 3 (Two-bottle preference test). (C) Comparison of behavioral data [9] and simulation data in control (all-satisfied) and water/sodium-depleted group. Control groups show minor intakes, and both depleted groups demonstrate copious volumes of intake in both sets of data. (D) In the Na-depleted group, water intakes are slight, and saltwater intake is increased. (E) The water-deficient groups show abundant water intakes and non-negligible saltwater intakes (F-I) Transitions of the HRL models (F) The water/sodium-deficient HRL model shows strong increases of values of water and saltwater intake. (G) The action value of sodium intake singly soars in the Na-depleted model. (H) The water-depleted model shows increased values of water and saltwater intake. (I) The control model refuses both intakes.

The results of the simulations revealed that the control group with fulfilled initial internal states showed continuously decreased action values of both water intake and salt intake, and the individuals in this group mostly chose to perform the do-nothing behavior. As a result, both internal states remained flat in the ideal state (Fig 3I). Accumulating these intakes, consumption from water bottles and saltwater bottles was low-keyed (Fig 3C-3E). In contrast, in the water/salt-depleted group, the action values of both water intake and salt intake rapidly increased. Reflecting these increases, both the water and sodium states also increased (Fig 3F). The accumulation of these consumptions was evidently larger in the depleted than in the control group (Fig 3C). In other circumstances, the sodium-depleted group, which was provided with large amounts of water, showed an increased action value of salt intake, resulting in an increased sodium state (Fig 3G). The total intake of water was slight, whereas salt intake was dominant (Fig 3D). The water-deficient group exhibited notable behaviors. First, both the action values of water intake and salt intake increased. The action value of water intake was larger than that of salt intake. Saltwater intake slightly increased (Fig 3H). These trends were similar to those observed in the actual animals assessed using the two-bottle preference test [9] (Fig 3C).

### Experiment-4: Simulated chemogenetic neural manipulation in the salt appetite network

Among the brain regions related to sodium appetite, it is assumed that neurons in the lateral parabrachial nucleus (LPBN^Htr2c^ neurons) have a suppressive function toward sodium appetite because artificial inhibition of that neuron with chemogenetic neural manipulation (designer receptors exclusively activated by designer drugs/DREADD) increases sodium appetite [22]. LPBN^Htr2c^ neurons project to the central amygdala (CeA). Based on these previous findings, in this experiment, we hypothesized that LPBN^Htr2c^ neurons have the function to inflict a negative bias on the action value of sodium appetitive behavior. The hypothesis was implemented as an account of action values in the HRL model, as follows:

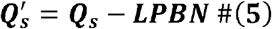

where *Q’* is the value of an action selection function. Additionally, the inhibition of LPBN^Htr2c^ neurons by DREADD was represented by the cancelation of the negative bias (i.e., LPBN in equation 5 was set to 0). With this implementation, we tested our hypothesis about LPBN neurons by comparing the simulation and the observation in an actual animal experiment (Fig 4A) [22].

**Fig 4.**
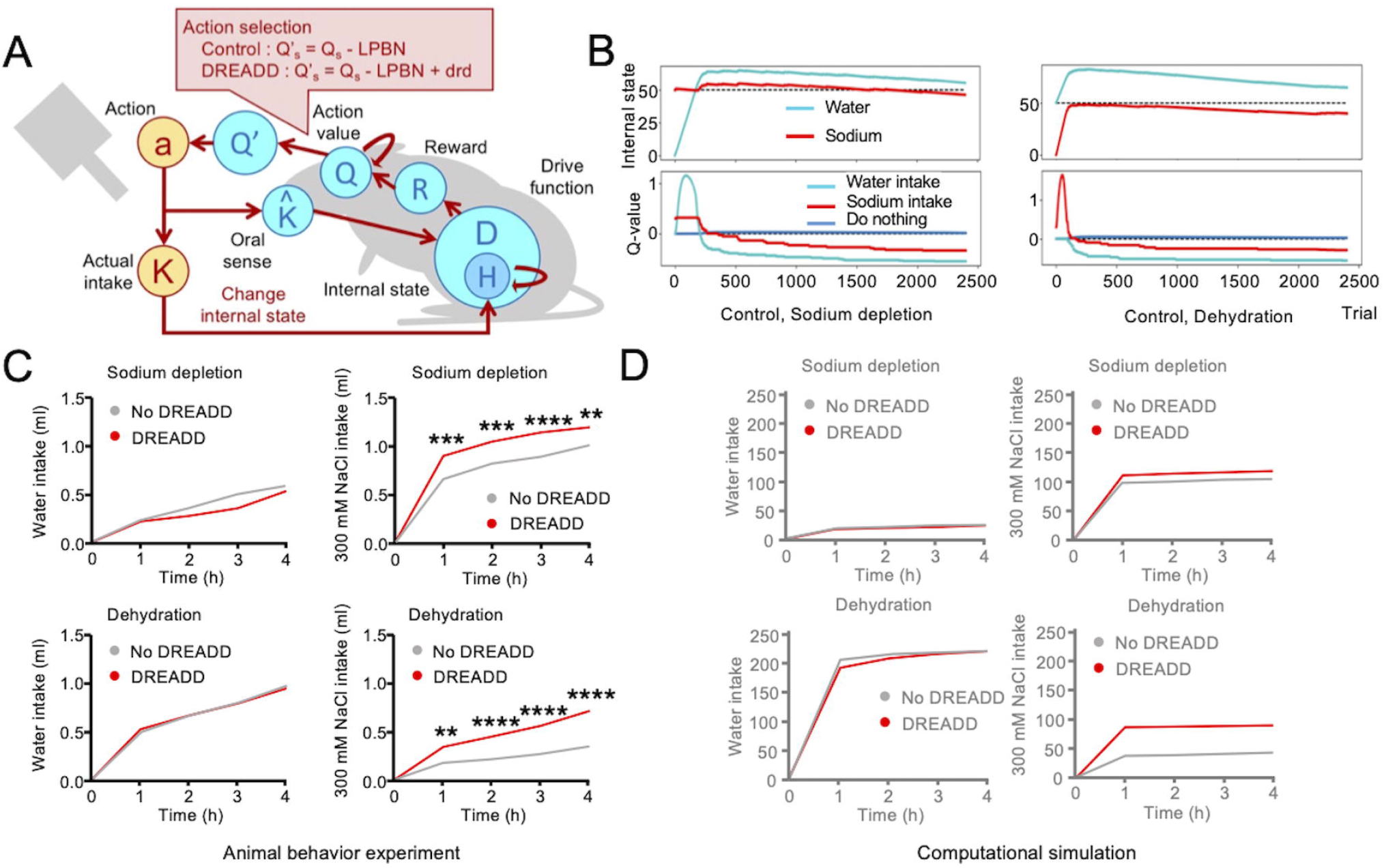
Simulation of designer receptors exclusively activated by designer drug (DREADD), which inhibits tonic neuronal suppression of sodium appetite. (A) Schematic illustration of experiment 4: DREADD (B) Transitions of control (no-DREADD) models. Sodium intake values were subtracted from these plots by a negative bias toward sodium appetite. (C) DREADD experiments increased sodium intake [22]. (D) Homeostatic reinforcement models demonstrate the equivalent behaviors.

This experiment was performed in a two-bottle preference test using water and saltwater [22]. There were four groups of mice based on the depletion of water/salt and application of DREADD, namely water-depleted/control (no DREADD), sodium-depleted/control, water-depleted/DREADD, and sodium-depleted/DREADD.

Both DREADD groups showed more saltwater intake than the control group. However, no clear differences were observed between the no DREADD groups. In the sodium depletion groups, water intake was slight in both the no DREADD and DREADD groups. In both groups, saltwater intake rapidly increased. The intake of the DREADD group was higher than that of the no DREADD group. Water intake increased sharply in the dehydration groups. There was no clear difference between the DREADD and no DREADD groups. The control (no-DREADD) model showed maintained homeostasis (Fig 4B). Saltwater intake of the dehydration groups was slight, but the DREADD group showed a much higher intake than that of the no DREADD group. These trends successfully replicated those observed in the actual animal experiments (Fig 4C and 4D).

## Discussion

In this study, we attempted to provide a mechanistic understanding of sodium appetite behavior using the HRL model. In the simple simulation of experiment 1, we confirmed that the HRL model successfully reproduced homeostasis-like behaviors by regulating sodium appetite, (i.e., approach and avoidance behavior to sodium). In addition, based on the assumption that the sense of taste is a predictor of changes in internal states, the HRL model successfully reproduced the previous observations of the intragastric infusion test that cannot be explained by the classical drive reduction theory [19]. These results support the idea that sodium appetitive behavior can be understood as a reinforcement learning process.

This idea is consistent with previous findings that the reward learning system is involved in sodium appetite behaviors. For instance, the activity of dopaminergic neurons in the VTA, which is thought to have a robust relationship with the reinforcement learning process in the brain [23, 24], was increased when sodium-depleted mice licked saltwater. In contrast, pharmacological inactivation of neural projections from the VTA to the nucleus accumbens decreased sodium intake [18]. In addition, optogenetic excitation of VTA dopaminergic neurons suppresses sodium intake in sodium-depleted mice [17]. The HRL model may provide insight into integrating these previous findings.

In the HRL model, the sense of taste was hypothesized to predict changes in the internal state. The latter means that the salty stimulus represents an immediate inducer of reinforcement, but this was not the case for the actual changes in the internal state. Nevertheless, only actual intake may result in the satiation of internal states. This assumption is consistent with the fact that gastric infusion of water does not work as a reinforcer, in contrast to oral intake of water, which can work as a reinforcer [2, 25]. In addition, artificial sweeteners, including saccharine and sucralose, can function as reinforcers [26-28], although their effects are relatively weaker than those of sucrose, which induces substantial changes in the internal state. From the perspective of computational theory, this assumption of the HRL model corresponds to the predictive processing (also referred to as predictive coding, or active inference) theory, in the sense that homeostasis is understood as the prediction of interoceptive sensory states and minimization of prediction error [29]. As such, homeostasis and sodium appetite behavior may provide an ideal research field to unify these computational theories.

In experiment 3, we extended the HRL model to multi-modal data, and successfully reproduced the behavioral tests in which water and sodium appetite regulate each other. In the current study, as the simplest attempt, the internal states of sodium and water were assumed to contribute equally. However, in an actual biological system, the homeostatic maintenance of water and sodium may not be exactly equal. For example, the effects of intragastric infusion of water and saltwater on the appetite of water and sodium may have different timescales [9, 11]. A more detailed implementation of such differences in water and sodium appetite may provide novel insights for understanding the system-level mechanisms of sodium appetite.

Finally, we successfully replicated the characteristic features of LPBN^Htr2c^ suppression experiments using DREADD. In the proposed model, we assumed that LPBN^Htr2c^ neurons provide a tonic negative bias toward sodium appetite via the central amygdala. That is, the LPBN-amygdala projection provides a negative bias of action selections and has no direct contribution to the learning of the Q-value. Based on this assumption, the learning of the Q-value canceled this negative bias, resulting in the inhibition of LPBN neurons, enhancing the relative value of salt intake, and provoking a rapid increase in salt-intake behavior. This assumption is consistent with previous findings that an immediate increase in sodium craving by sodium depletion may not be mediated by learning processes [30]. In addition, CeA is an essential region for both hedonic and aversive intakes. For example, CeA prepronociceptin-expressing neurons are activated by hedonic intake and promote palatable food consumption [31]. In contrast, activation of PKC-d+ neurons in the lateral subdivision of CeA inhibits feeding [32]. Further investigation of the neural connections from LPBN to CeA, together with the HRL model, may provide fundamental information for the development of more precise algorithms.

Here, we discuss the significance of constructing a computational model for sodium appetite. To understand complex systems such as the brain, investigations from three levels are essential, namely computational theory, representation and algorithm, and hardware implementation [33]. In this study, we provided a mechanistic explanation of sodium appetite behavior by bridging previous findings from the aspects of these three levels. Although the model behaviors were evaluated only for their quantitative similarities with the actual animal experiments, the model can also provide quantitative predictions of unobservable latent variables, such as reward prediction error, values of actions (motivation toward nutrient intake), and prediction of internal states. Investigations of neural correlates of such latent variables may provide a deeper understanding of the underlying neural mechanisms of sodium appetite and homeostatic behavior.

## Methods

### Sodium homeostasis

An overview of the HRL model is described in the Results section (see Equations 1–4). To investigate the applicability of the HRL model to sodium appetite behavior, we performed a sodium intake test (Experiment 1). The computation algorithm is illustrated in Fig 1A. In this experiment, only the internal state of sodium was considered. An external state (*S*_*0*_) and two actions, do nothing (*a*_*0*_) and intake (*a*_*1*_), were assessed (Fig 1B). Action selection depends on the relative magnitudes of the values of each action (Q-value), following the soft-max function:

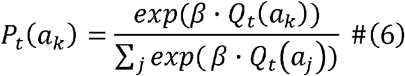

where *P*_*t*_*(a*_*k*_) is the probability of an action *a*_*k*_ to be selected at time *t, and* β is the inverse temperature, a parameter controlling the randomness of an action. In Experiment 1, the Q-values of both actions were set to 0. Therefore, the first action was randomly chosen. When intake behavior was performed, the internal sodium state increased with *K*, a constant parameter defining the size of sodium intake. When nothing was chosen, *K* was set to 0. At *t* = 0, to represent sodium depletion, the first internal state (*H* = 0) was far lower than the ideal state (H*=50). At this stage, the value of the drive function was large because the drive function corresponds to a type of distance from the internal state of time t (*Ht*) to the ideal state (H* = 50) (Equation 1). If an agent performed the intake behavior at this moment, the internal state increased and the drive function became smaller, resulting in a positive reward (Equation 2).

In addition, the natural decrease of sodium state was implemented as follows using the temporal decay constant *τ* (Equation 7).

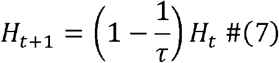

As a result, the calculation of the reward value was determined as follows (Equation 8).

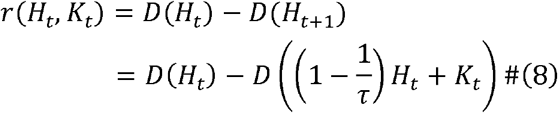

To renew Q-values with the reward value, we used a conventional Rescorla-Wagner model [35], where *i* indicates each action, *⍰* is learning rate, and *r - Q* represents reward prediction error [35]. After this update of the Q-values, the agent chooses the next action. The detailed values of the parameters of this simulation are listed in Table S1.

### Oral sense as a predictor of changes in internal states

In experiments 2–4, we introduced oral sense *K^*, which represents the prediction of changes in the internal state. The definition of the reward (Equation 3) was also updated as follows:

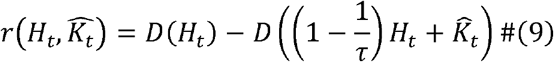

Thus, the reward was defined as changes in internal states and taste sensory input as predictors for changes in internal states. The other functions were the same as those in experiment 1.

The behavioral experiment of intragastric infusion is shown in Fig. 2A. The animals were grouped into three groups: control, intragastric, and oral stimulation. Only the control group was salt-depleted. In the intragastric (IG) infusion group was sodium depleted. In addition, an intragastric cannula was inserted into the gut of the mice. Before the intake test, saltwater was directly infused into the gut. In the oral stimulation group, the animals were stimulated with a strong salty stimulus in the intake test. To simulate the IG infusion group, we hypothesized that the internal state of sodium was set to the level of half-satisfaction at the beginning of the intake test. To model the oral stimulation group, a salty taste was supplied, regardless of the selected actions. The definitions of states and actions were the same as in Experiment 1 (Fig 1B). The algorithm including taste sensory input as a predictor of changes in the internal state is illustrated in Fig 2A. The detailed values of the parameters used in this experiment are shown in Table (in Table S2).

### Two-bottle preference test

For the simulation of the two-bottle preference test in experiment-3, we set two internal states corresponding to water and sodium. Thus, in Equation 1, the number of dimensions of internal state *N* was set to 2. Each internal state updates as follows:

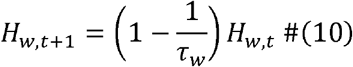

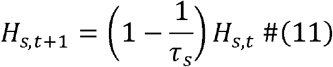

where *H*_*w*_ is water state, and *H*_*s*_ is sodium state.

The detailed parameters were partially different from those of the taste expectation. The detailed values of the parameters in experiment 3 are shown in Table S3.

### DREADD experiment

To simulate the DREADD experiment, we assumed that LPBN^Htr2c^ neurons provided tonic suppression of sodium appetite as implemented by a tonic negative bias in the selection of sodium intake action as follows:

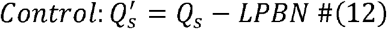

where LPBN is a positive constant value corresponding to the negative bias of the LPBN^Htr2c^ neuron. Note that Q’ is used only for action selections and Q-values are updated with previous Q-values and the reward, regardless of Q’ and the tonic bias (Equation 4).

DREADD treatment was implemented as the cancelation of this negative bias by adding the positive value of *drd* as follows:

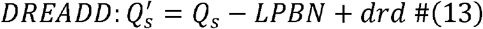

In the current experiment, the values of LPBN and *drd* were set to be the same. At the beginning of the saltwater intake test, the initial Q-value for salt intake behavior was set to LPBN, corresponding to the state, where *Q’* for both actions was 0 (i.e., the probabilities for salt intake and do-nothing were 0.5). The initial water state was set to 0 for the dehydration group, and the initial sodium state was set to 0 for the sodium-depleted group. The detailed values of the parameters in Experiment 3 are shown in Table S4.

## Supporting information captions

**S1 Table. Free parameters for experiment 1**.

Parameters for experiment of sodium homeostasis according to the HRL.

**S2 Table. Free parameters for experiment 2**.

Parameters for a simulation with sense of taste as a predictor of changes in internal states.

**S3 Table. Free parameters for experiment 3**

Parameters used for multi-dimensional HRL.

**S4 Table. Free parameters for experiment 4**.

Parameters for a simulation of chemogenetic neural manipulation

