## Supplemental Table 2 for "Computational mechanisms of osmoregulation: a reinforcement learning model for sodium appetite"

**S2 Table. Free parameters for experiment 2**

| Parameter | Value | Contents |
| --- | --- | --- |
| $\boldsymbol{\alpha}_{\boldsymbol{1}}$ | 0.006 | Learning rate of Q-values |
| $\boldsymbol{\alpha}_{\boldsymbol{2}}$ | 0.6 | Learning rate from taste sensory input to change of internal state |
| $\boldsymbol{\beta}$ | 1.67 | Inverse temperature of soft-max function |
| $\boldsymbol{\tau}$ | 100 | Attenuation rate of internal state |
| $\boldsymbol{m}$ | 3 | Free parameter of the drive function |
| $\boldsymbol{n}$ | 4 | Free parameter of the drive function |
| $\boldsymbol{H}^{\boldsymbol{*}}$ | 50 | Ideal internal state |
| $\boldsymbol{K}$ | 1 | Sodium intake (per lick) |
| $\hat{\boldsymbol{K}}$ | 1 | Taste sensory input (per lick) |
