## Supplemental Table 3 for "Computational mechanisms of osmoregulation: a reinforcement learning model for sodium appetite"

**S3 Table. Free parameters for experiment 3**

| Parameter | Value | Contents |
| --- | --- | --- |
| $\boldsymbol{\alpha}_{\boldsymbol{1}}$ | 0.03 | Learning rate of Q-values |
| $\boldsymbol{\alpha}_{\boldsymbol{2}}$ | 0.3 | Learning rate from taste sensory input to change in the internal state |
| $\boldsymbol{\beta}$ | 10 | Inverse temperature of soft-max function |
| $\boldsymbol{\tau}_{\boldsymbol{w}}$ | 7000 | Attenuation rate of water state |
| $\boldsymbol{\tau}_{\boldsymbol{s}}$ | 7000 | Attenuation rate of sodium state |
| $\boldsymbol{m}$ | 3 | Free parameter of the drive function |
| $\boldsymbol{n}$ | 4 | Free parameter of the drive function |
| $\boldsymbol{H}^{\boldsymbol{*}}$ | 50 | Ideal internal states |
| $\boldsymbol{K}_{\boldsymbol{w}}$ | 0.3 | Water intake (per lick) |
| $\boldsymbol{K}_{\boldsymbol{s}}$ | 0.5 | Sodium intake (per lick) |
| $\hat{\boldsymbol{K}_{\boldsymbol{w}}}$ | 0.3 | Taste sensory input of water (per lick) |
| $\hat{\boldsymbol{K}_{\boldsymbol{s}}}$ | 0.5 | Taste sensory input of sodium (per lick) |
